# Impact of baseline culture conditions of mouse-derived cancer organoids when determining therapeutic response and tumor heterogeneity

**DOI:** 10.1101/2021.10.12.464087

**Authors:** Rebecca A. DeStefanis, Autumn M. Olson, Alyssa K. DeZeeuw, Amani A. Gillette, Gioia C. Sha, Katherine A. Johnson, Jeremy D. Kratz, Cheri A. Pasch, Linda Clipson, Melissa C. Skala, Dustin A. Deming

## Abstract

Representative models are needed to screen new therapies for patients with cancer. Cancer organoids are a leap forward as a culture model that faithfully represents the disease. Mouse-derived cancer organoids (MDCOs) are becoming increasingly popular, however there has yet to be a standardized method to assess therapeutic response and identify subpopulation heterogeneity. There are multiple factors unique to organoid culture that could affect how therapeutic response and MDCO heterogeneity are assessed. Here we describe an analysis of nearly 3,500 individual MDCOs where individual organoid morphologic tracking was performed. Change in MDCO diameter was assessed in the presence of control media or targeted therapies. Individual organoid tracking was identified to be more sensitive to treatment response than well-level assessment. The impact of different generations of mice of the same genotype, different regions of the colon, and organoid specific characteristics including baseline size, passage number, plating density, and location within the matrix were examined. Only the starting size of the MDCO altered the subsequent growth. Here we establish organoid culture parameters for individual organoid morphologic tracking to determine therapeutic response and growth/response heterogeneity for translational studies using murine colorectal cancer organoids.

## Introduction

Colorectal cancer (CRC) is the second leading cause of cancer related deaths in the United States and is estimated to cause approximately 53,000 deaths in 2021 [1, 2]. Clinical treatments for metastatic CRC have shifted drastically over the past decade owing to a greater understanding of how the molecular profile of a cancer can guide clinical care strategies. More specifically, precision-guided approaches, such as anti-epidermal growth factor receptor inhibitors and immune checkpoint inhibitors are used for *KRAS/NRAS/BRAF* wild-type and mismatch repair deficient CRCs, respectively [3]. Despite these clinical advancements, preclinical models available to identify and study potential therapeutic strategies for these and other emerging subtypes of CRC remain few. Further, models such as historical 2-dimensional cell culture are limited in their ability to faithfully represent the disease [4-6].

Cancer organoid cultures continue to be a major advance for studying therapeutic strategies. Organoid cultures are three-dimensional (3D) cell cultures that can be isolated from patient or mouse tumors and are grown in extracellular matrices, such as Matrigel or collagen. Compared to traditional 2D immortalized cell lines, cancer organoids better recapitulate the tumor from which they were derived, both morphologically and molecularly [7-21]. Historically, it has been difficult to establish 2D cell lines from more common cancer types in part owing to their inability to adhere to plastic [22, 23]. The development of cancer organoids has significantly increased our ability to establish patient specific cultures across cancer types.

With this advancement in culture techniques, research groups have utilized both mouse and patient-derived cancer organoids to test pre-clinical hypothesis-driven combination therapies and to identify novel therapeutic strategies with high-throughput drug screens [8, 24-28]. Given their 3D structure, traditional therapeutic response assessments used for 2D cultures are not directly transferable to organoid cultures. For this reason, multiple methods have been developed or adapted to assess response in these cancer organoids. Most assays measure metabolic activity of the cells at the well-level (*e*.*g*., CellTiter-Glo). One major pitfall to this method is its inability to evaluate the heterogeneity within a culture because these assays measure the gross response of the population within a given well. Additionally, these types of assays are not suitable to longitudinal monitoring, and thus, are very sensitive to baseline plating of the organoids, which is more challenging to control than with 2D cultures. To address this, our group has developed assays that measure the change in size or diameter of individual organoids over the course of the study. This method allows for the examination of an individual response of an organoid in addition to a population response [10, 18, 19, 29-33].

With the goal of using these organoid cultures as preclinical models to identify and confirm new therapeutic strategies, it is important to understand whether certain culture conditions affect growth and therapeutic response. Besides studies of media and supplements, limited data exists regarding the effect of baseline culture conditions on growth and response in these heterogeneous organoid cultures [26, 29, 34]. Several groups have investigated how different factors added to the media affect the maintenance and development of different organoid models [26, 27, 35-39]. However, no prior studies have performed a comprehensive analysis to examine how factors other than the media conditions alter growth or response of organoids. Here, we have evaluated a mouse-derived cancer organoid (MDCO) model developed by our group to address this knowledge gap. These MDCOs were established and analyzed over a 5-year period, enabling a direct comparison of data collected years apart from independent cultures. Specifically, we investigated if significant variation was seen between MDCOs derived from different mice of the same genotype or regions of the colon. Additionally, we assessed culture conditions including baseline size or location within the Matrigel droplet of individual MDCOs, the passage number of the line, and the density of the culture to determine whether these baseline conditions affect growth and response (Table 1).

**Table 1.** Number of individual organoids used in the growth and response organoid characteristic analyses. The number (N) of organoids used in each organoid characteristic analysis is listed, along with the corresponding figure number. For the response analyses, all PI3K pathway inhibitors used are listed.

| Organoid characteristics | Inhibitors assessed | All organoids |  | Organoids with CPA applied |  |
| --- | --- | --- | --- | --- | --- |
|  |  | N of organoids | Figure | N of organoids | Figure |
| Growth |  |  |  |  |  |
| Change point analysis | NA | 994 | 2 | NA | NA |
| Mouse | NA | 1001 | 3a | 834 | S1a |
| Tumor location | NA | 855 | 3b | 734 | S1b |
| Passage number | NA | 968 | 4a | 817 | S2a |
| Location within Matrigel droplet | NA | 543 | 5b | 493 | S3a |
| Density | NA | 3580 | 6a | 3277 | S4a |
| Density (prospective) | NA | 362 | 6d | NA | NA |
| Response |  |  |  |  |  |
| Mouse | AZD2014, BYL719, copanlisib, everolimus, NVP-BEZ235, TAK-228 | 2480 | 3a | 2222 | S1c |
| Tumor location | AZD2014, BYL719, copanlisib, everolimus, NVP-BEZ235, TAK-228 | 2203 | 3b | 2002 | S1d |
| Passage number | AZD2014, BYL719, copanlisib, everolimus, NVP-BEZ235, TAK-228 | 2380 | 4a | 2136 | S2b |
| Location within Matrigel droplet | AZD2014, BYL719, copanlisib, everolimus, NVP-BEZ235, TAK-228 | 1292 | 5b | 1186 | S3b |
| Density | AZD2014, BYL719, copanlisib, everolimus, NVP-BEZ235, TAK-228 | 6170 | 6a | 6021 | S4b |
| Density (prospective) | copanlisib | 498 | 6d | NA | NA |

## Results

### Cancer organoids grow heterogeneously within a culture

We have previously shown the use of MDCOs derived from *Fc*^*1*^*Apc*^*fl/+*^ *Pik3ca*^*H1047R*^ (APPK) transgenic mouse CRCs as a model to examine potential therapeutic strategies, specifically those targeting the PI3K pathway [40]. Here we examine MDCOs from this model to assess the effects of baseline culture conditions on growth and response to targeted therapies. Briefly, these MDCOs are cultured in the extracellular matrix Matrigel and plated in droplets with media overlayed on top (Fig 1a). The diameter of each organoid was measured at baseline and after 48 hours with care taken to track the change in growth of each individual organoid over time. Within each culture, differential organoid growth is observed (Fig 1c). This heterogeneity is maintained across numerous cultures derived from different mice and yields similar distributions of the change in organoid diameter (Fig 1b).

**Figure 1.**
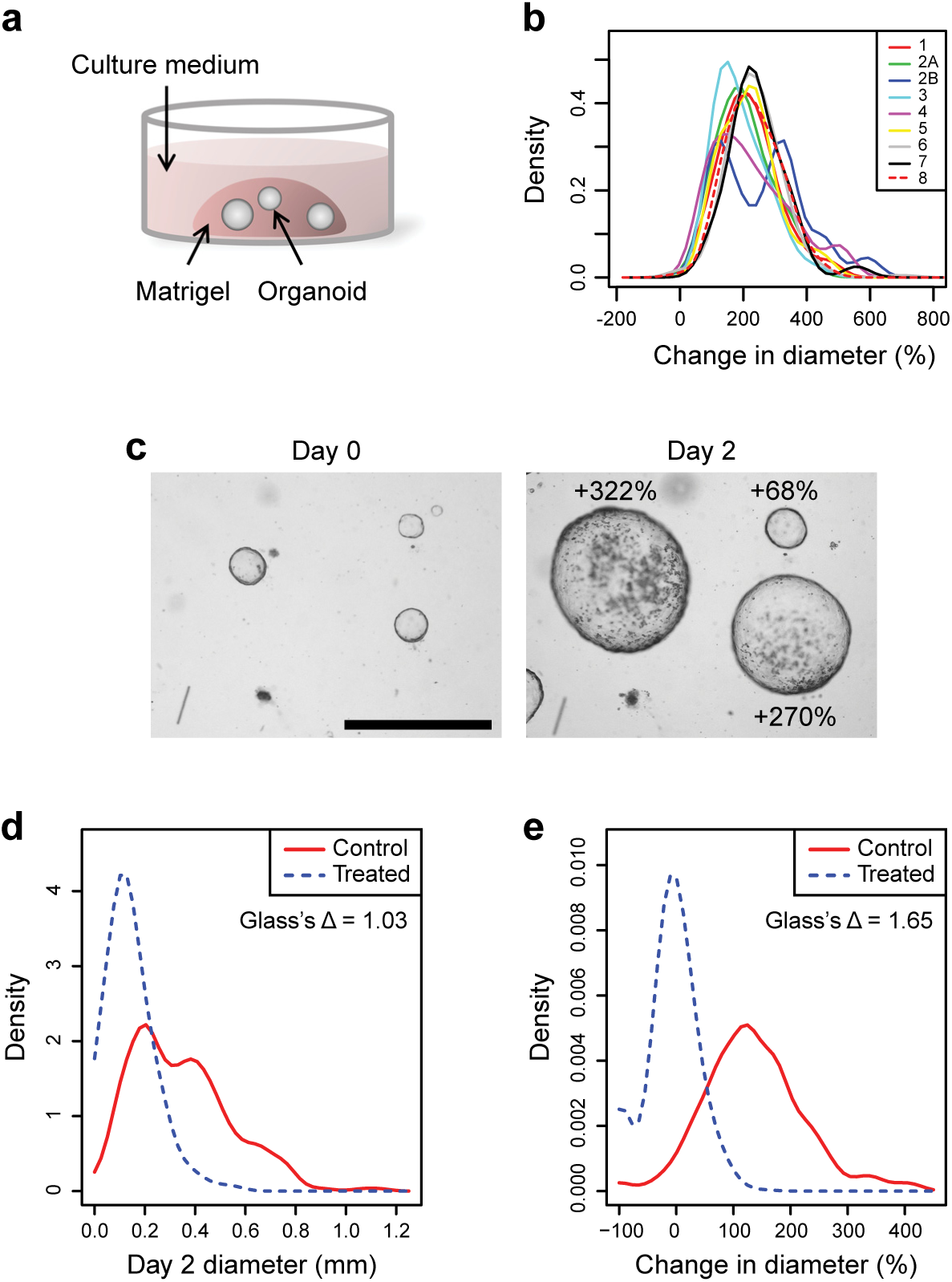
Mouse derived cancer organoids grow heterogeneously within a culture. (a) Graphic illustrating how MDCOs isolated from Fc1 Apcfl/+ Pik3caH1047R mice are cultured within a Matrigel matrix, with feeding media overlayed. For therapeutic studies, overlayed feeding media is replaced with new media containing drug. (b) Kernel density plot comparing the growth of MDCOs at the population level derived from 8 different mice of the same genotype. Each line indicates the growth of MDCOs in feeding media derived from one mouse. Note that mouse 2 had two MDCO lines derived from two distinct tumors (line 2A and 2B). (c) Representative image of the heterogeneous growth rates of MDCOs within a culture over 48 hours. (d) Density plot comparing the growth and response of MDCOs using only the final diameter (mm) after 48 hours. This analysis represents a well level analysis where MDCOs are only evaluated on the final day of a study. Effect size was calculated using Glass’s delta to compare the treatment group to the control group. (e) Kernel density plot comparing the growth and response of MDCOs using their percent change in diameter over the 48 hour incubation. This analysis examines the MDCOs on an individual level to determine how individual MDCOs change in diameter over the course of a study. Effect size was calculated using Glass’s delta to compare the treatment group to the control group. The studies used in (d) and (e) include MDCOs treated with normal feeding media (Control) or copanlisib (200nmol/L) (Treated) for 48 hours. Representative images from (c) are shown at the same magnification. Size bar, 1 mm.

Multiple methods have been developed to assess therapeutic response using 3D organoids. Most of those methods examine the organoids on a whole-well basis to examine the population response, usually on the final day of analysis. Metabolic assays are grossly affected by organoid plating which is much more challenging to control than in classic 2D cultures. Even in the setting where each individual organoid diameter is examined, the 48-hour time point measurements alone lack sensitivity to detect treatment response (Fig 1d) compared to changes in diameter of individual organoids over 48 hours, likely due to differences in baseline organoid sizes (Fig 1e). Additionally, a larger effect size is observed between the treatment and control for the change in diameter versus the 48-hour time point alone (Glass’s delta values 1.65 vs 1.03, respectively) [41]. This ability to examine response on the single organoid level over time results in greater sensitivity to treatment response. To examine whether any of the heterogeneity observed could be due to organoid culture conditions rather than individual organoid biology, further investigations into how changes in the culture conditions could alter organoid growth were performed.

### Growth rate of MDCOs change as a function of their starting size

Previously, using a standard change point analysis (CPA), we reported that APPK MDCOs <373 µm at baseline had a similar growth rate. However, MDCOs that were ≥373 µm had a reduction in growth rate and were therefore excluded from analysis [40]. We reapplied this change point analysis to include all of the control MDCOs in this analysis (n =1019), which include those from our previously published work, and found that with this larger dataset, 308 µm is the more accurate change point value. This indicates that MDCOs that are ≥308 µm at baseline should be excluded from downstream analyses because the growth rate changes as a function of their size. (Fig 2) We applied this cutoff to our analyses and found that <450 individual MDCOs were ≥308 µm at the beginning of these studies (∼12% of the total population). Approximately 40% of the MDCOs were controls and 60% were MDCOs treated with therapeutic agents. MDCOs with baseline sizes above the change point were excluded from subsequent supplemental analyses. (Suppl Fig 1-5).

**Figure 2.**
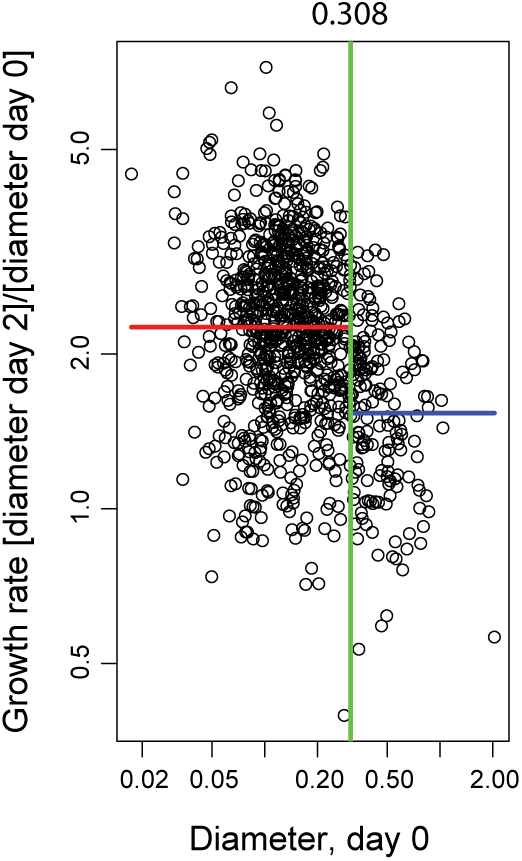
A standard change point analysis demonstrates that the growth rates of the MDCOs vary as a function of their size. A change point analysis at 308 μm was determined using all control APPK MDCOs. When the change point analysis was applied to all studies, only MDCOs <308 μm on day 0 were used in the analyses (Suppl Figs 1-4). The green line indicates the calculated change point value. The red line indicates the geometric mean of the data points that are lower than the determined change point value. The blue line indicates the geometric mean of the data points that are greater than the determined change point value. (n = 994)

### MDCOs derived from different mice of the same genotype and from different regions of the colon do not vary significantly in growth or drug response

One important feature of transgenic mouse models is the ability to generate mice within litters and across generations that have identical activation of transgenes and near identical genetic backgrounds. For this reason, we can isolate new or additional APPK MDCO lines from different mice of the same genotype both within a litter and across generations. To confirm that MDCO lines from different mice of the same genotype have similar growth distributions, a total of 8 different MDCO lines from 8 different cancers were isolated from 7 mice, two lines being isolated from two different tumors in one mouse. Minimal variation was seen across the growth of these MDCO lines as the majority of the individual MDCOs were within 1 standard deviation (SD) of the population mean (PM) (125 ± 88%) (Fig 3a).

**Figure 3.**
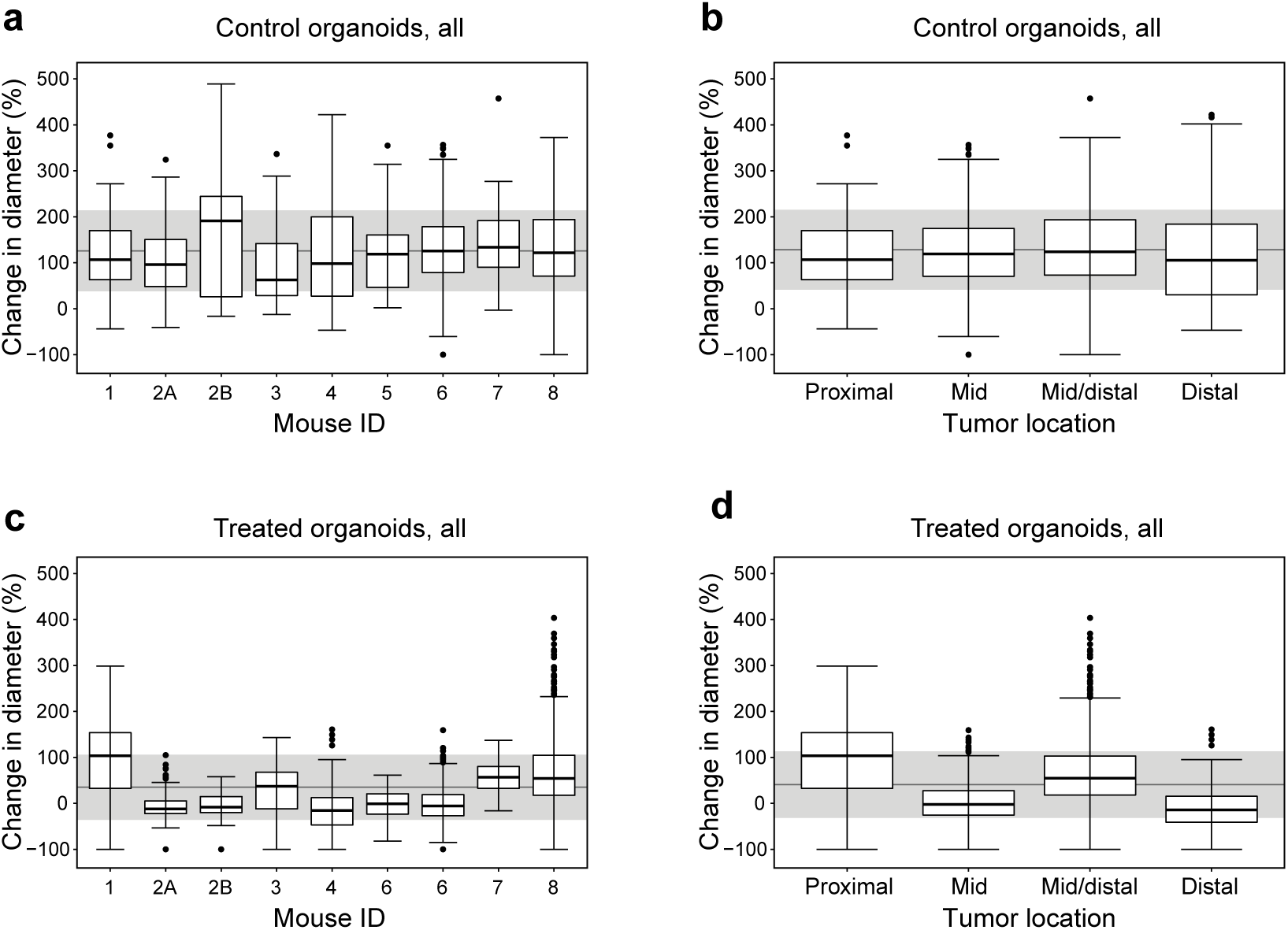
MDCOs derived from different mice of the same genotype or from different regions of the colon do not vary significantly in growth or drug response. Box and whisker plots displaying control organoids from (a) different mice of the same genotype and (b) tumors isolated from different regions of the colon (n = 1001and n = 855, respectively). Box and whisker plots displaying treated organoids (c) derived from different mice of the same genotype and (d) tumors isolated from different regions of the colon (n = 2480 and n = 2203, respectively). Each number represents a different mouse. Note that the MDCO line derived from mouse 1 was isolated from a proximal colon tumor and mouse 2 had two MDCO lines derived from two different colon tumors. Closest to the small intestine is the proximal colon moving more distally to the mid colon and finally the distal colon. Note the proximal colon has a different gross histology than the mid or distal colon. In all box and whisker plots the grey line indicates the population change in diameter mean while the grey shading indicates one standard deviation above and below the population mean.

The *Fabp1-Cre* drives Cre-recombinase expression and subsequent constitutive recombination of transgenes in the epithelial portion of the distal small intestine and the large intestine. Therefore, tumor formation can occur anywhere in those regions of recombination. Tumors are isolated from multiple regions of the large intestine to establish APPK MDCO lines. APPK MDCO lines were derived from tumors in the proximal colon, which is closest to the stomach, the mid colon, mid/distal colon, and distal colon. It is important to note that the gross histology of the proximal colon is different from that of the mid and distal colon. Additionally, due to this difference in gross histology, only one mouse line (mouse 1) was found to have been isolated from the proximal colon. No significant variation was observed in the growth of the MDCO lines due to their original tumor location as most individual organoids were within 1SD of the PM (127, ± 87%) (Fig 3b). MDCOs that were ≥308 µm at baseline were then removed from these analyses, based on the CPA from Fig 2. We still found that growth was not affected by which mouse the line was derived from or which region of the colon the tumor was from as the majority of the MDCOs remained within 1SD of the PM (Mouse: 137 ± 88%; Tumor 138 ± 86%) (Suppl Fig 1a and 1b).

Both variation in the mouse from which the organoids were derived and original tumor location were evaluated as potential conditions that might alter treatment response, in this case to PI3K pathway inhibition. Work from our group, using APPK MDCOs, has demonstrated that dual MTORC1/2 inhibition is sufficient to induce a treatment response in *Pik3ca* mutant CRC. To assess therapeutic response, the overlayed medium was replaced with new medium containing drug, and the diameter of each organoid measured at baseline and after 48 hours to track how individual MDCOs and the population of MDCOs respond to a given therapy [40, 42]. These studies were used in this pooled analysis along with other studies that assessed the efficacy of PI3K pathway inhibitors. Altogether six PI3K pathway inhibitors with doses ranging from 5-500nmol/L including those that inhibit PI3K alone (BYL-719; α isomer), both PI3K/MTOR (BEZ-235, copanlisib), MTORC1 (everolimus), and both MTORC1/2 (AZD-2014, TAK-228) were used in this analysis (Table 2).

**Table 2.** PI3K pathway inhibitors and doses of inhibitors used in organoid response characteristics analyses. Multiple PI3K pathway inhibitors were assessed in the organoid response characteristic analyses ranging from 5-500nmol/L.

| Inhibitor | Target | Doses (nmol/L) |
| --- | --- | --- |
| AZD2014 | mTORC1/2 | 200, 300, 400, 500 |
| BYL719 | PI3Kα | 200 |
| Copanlisib (BAY 80-6946) | PI3K/mTOR | 5, 10, 100, 200, 400 |
| Everolimus | mTORC1 | 100, 200, 400 |
| NVP-BEZ235 | PI3K/mTOR | 100, 200, 400 |
| TAK-228 | mTORC1/2 | 100, 200, 400 |

The responses of all MDCOs treated with a PI3K pathway inhibitor were grouped based on the mouse number and the original tumor location. The mean population of treated MDCOs by mouse number and tumor location were 34% (± 70%) and 40% (± 72%), respectively. Interestingly a proportion of the individual MDCOs from mouse 1, which was the only MDCO line originally isolated from the proximal colon, fell outside of 1 SD from the PM both in the different mouse line assessment and original tumor location assessment. (Fig 3c and 3d) This could be due to the gross histological differences between the proximal colon and the rest of the colon, though more studies are needed to confirm this explanation. With the CPA applied, we observed similar results that indicated mouse number (37 ± 72%) and tumor location (42 ± 73%) do not affect the MDCO drug response (Suppl Fig 1c and 1d).

Overall, these data indicate that growth of MDCOs are not affected by being isolated from different mice of the same genotype or original tumor location. However, our studies suggest that MDCO lines isolated from the proximal colon versus mid, mid/distal, or distal colon could have some variation in how they respond to treatment. These differences suggest that MDCOs derived from the proximal colon should be further investigated and not directly compared to MDCOs derived from the mid, mid/distal, or distal colon.

### Passage number of cultures does not affect growth or drug response of APPK MDCOs

Once it was established that there was no significant variation in growth or response at the mouse level, we sought to look further in depth at specific baseline culture conditions. We next evaluated if passage number influences growth or response of the APPK MDCOs. It is well established that in traditional 2D cell lines higher passaged cells can develop significant alterations in growth rates, and in how they respond to therapies, among other characteristics [4-6, 43-45]. We analyzed MDCOs ranging from passage 1-15 across different lines from studies of PI3K pathway inhibitors. We observed that no significant variation was observed in growth as the majority of the untreated MDCOs were within 1SD of the PM (126 ± 89%) (Fig 4a). Similar observations with response were seen in the treated MDCOs. The response of the majority of the treated MDCOs fell within 1SD of the PM (36 ± 71%) (Fig 4b). We further excluded the MDCOs that were ≥308 µm as determined in Fig 2 and observed that the majority of MDCOs’ change in diameter fell within 1SD of the population mean for both growth (138 ± 88%) and response (38 ± 73%) (Suppl Fig 2). We did note at passage 5 that some of the MDCOs’ change in diameter fell below 1SD of the PM, however (Fig. 4a), the dataset was limited (n = 6 and n = 1 for growth and response assessment, respectively) making it difficult to accurately evaluate growth and response at this passage number.

**Figure 4.**
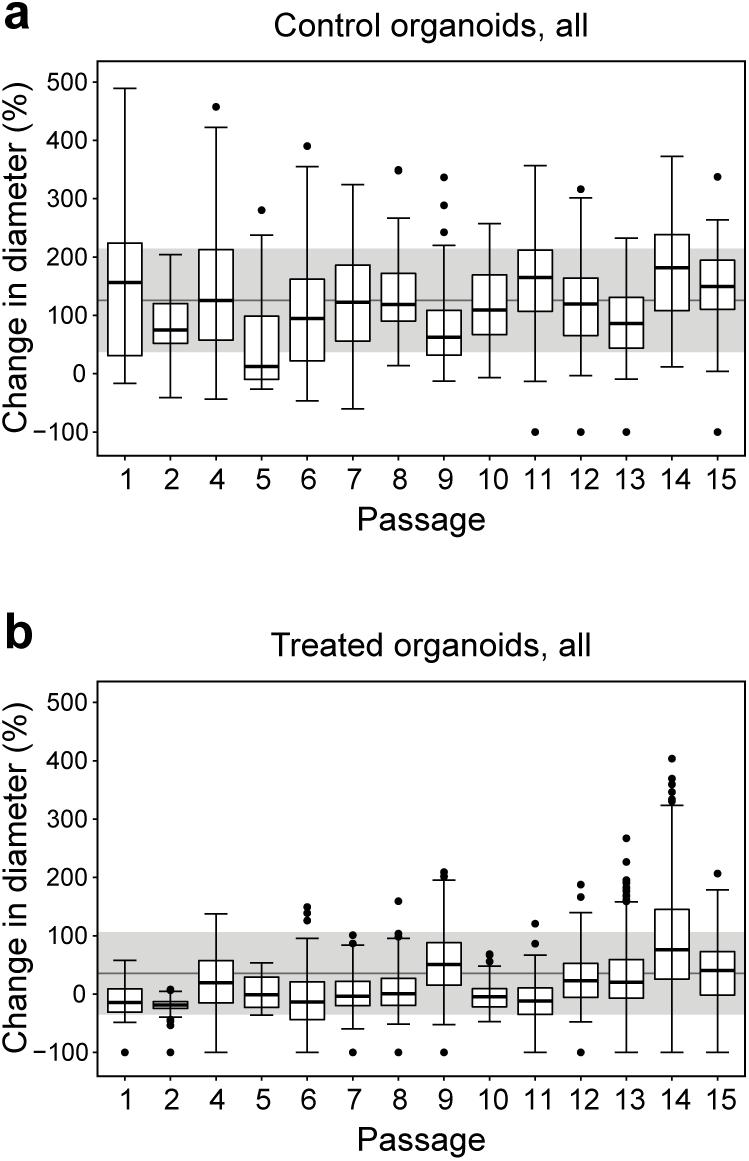
Passage number does not affect growth or drug response of APPK MDCOs. Box and whisker plots displaying the percent change in diameter for (a) growth and (b) drug response of MDCOs from passages 1-15 (n = 968 and n = 2380, respectively). In all box and whisker plots the grey line indicates the population change in diameter mean while the grey shading indicates one standard deviation above and below the population mean.

### Location or density of the APPK MDCO within the *in vitro* matrix does not affect growth or response

Unlike traditional 2D cultures, organoids are cultured within an extracellular matrix, such as Matrigel with feeding media overlayed (Fig 1a). To assess therapeutic response, the overlayed feeding media is exchanged with fresh media containing drug. Given this significant difference in culture methods we aimed to confirm that the MDCOs both at the periphery and towards the center of the Matrigel droplet were growing and responding to the same extent. To assess the location of each MDCO within the Matrigel droplet, the shortest distance from an individual MDCO to the edge of the Matrigel droplet, as seen in the field of view (FOV) was measured. This was plotted against each MDCO’s percent change in diameter (Fig 5a). The growth of the control MDCOs was not correlated with the location of the MDCOs within the Matrigel droplet (R^2^ = 0.0022) (Fig 5b). Similar observations were observed in the treatment response, as indicated by a low R^2^ value when comparing the treated MDCOs’ percent change in diameter to location within the Matrigel droplet (R^2^ = 0.0031) (Fig 5c). The change point analysis was then applied to these data. With the exclusion of the MDCOs that fell outside of the appropriate starting size (i.e. excluding organoids ≥308 µm) no correlation was observed between location of the MDCO and growth (R^2^ = 0.0014) or response (R^2^ = 0.0025) (Supp Fig 3).

**Figure 5.**
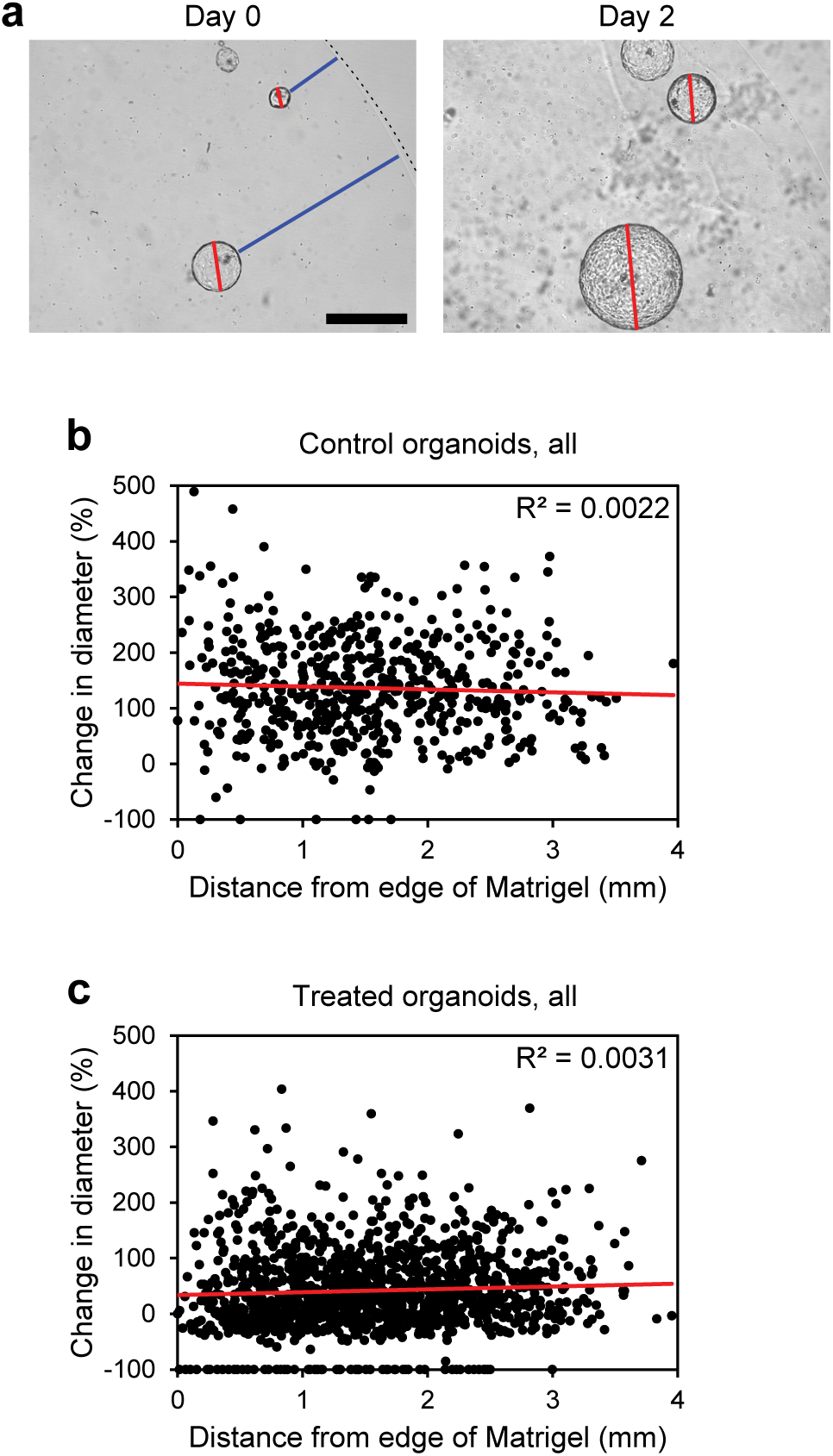
Location within the Matrigel matrix does not affect MDCO growth or drug response. (a) Representative images illustrating how location within the Matrigel matrix was done. Briefly, for each MDCO, the shortest distance (blue line) from the edge of the organoid to the edge of the Matrigel (dashed line) at the beginning of the study was measured using ImageJ. Scatter plots display individual organoids’ location within the Matrigel matrix vs their change in diameter in (b) growth and (c) drug response (n = 543 and n = 1292, respectively). Linear trend lines are indicated in red. The R2 values displayed in the top right corner indicate that no significant correlation exists between the MDCOs change in diameter and location within the Matrigel matrix. Representative images from (a) are shown at the same magnification. Size bar, 500 μm.

We next evaluated whether density, which is equivalent to confluency in traditional 2D cultures, affects the growth or response of the MDCOs. To do this, the total number of MDCOs/field of view was calculated and compared to the average percent change in diameter of the MDCOs within that field of view. No association was seen between the density and the growth or response of the MDCOs (R^2^ = 0.0252, R^2^ = 0.0823, respectively) (Fig 6a and 6b). Once the change point analysis was applied, similar observations of no correlation between density and growth or response were observed (R^2^ = 0.0041, R^2^ = 0.0644, respectively) (Supp Fig 4).

**Figure 6.**
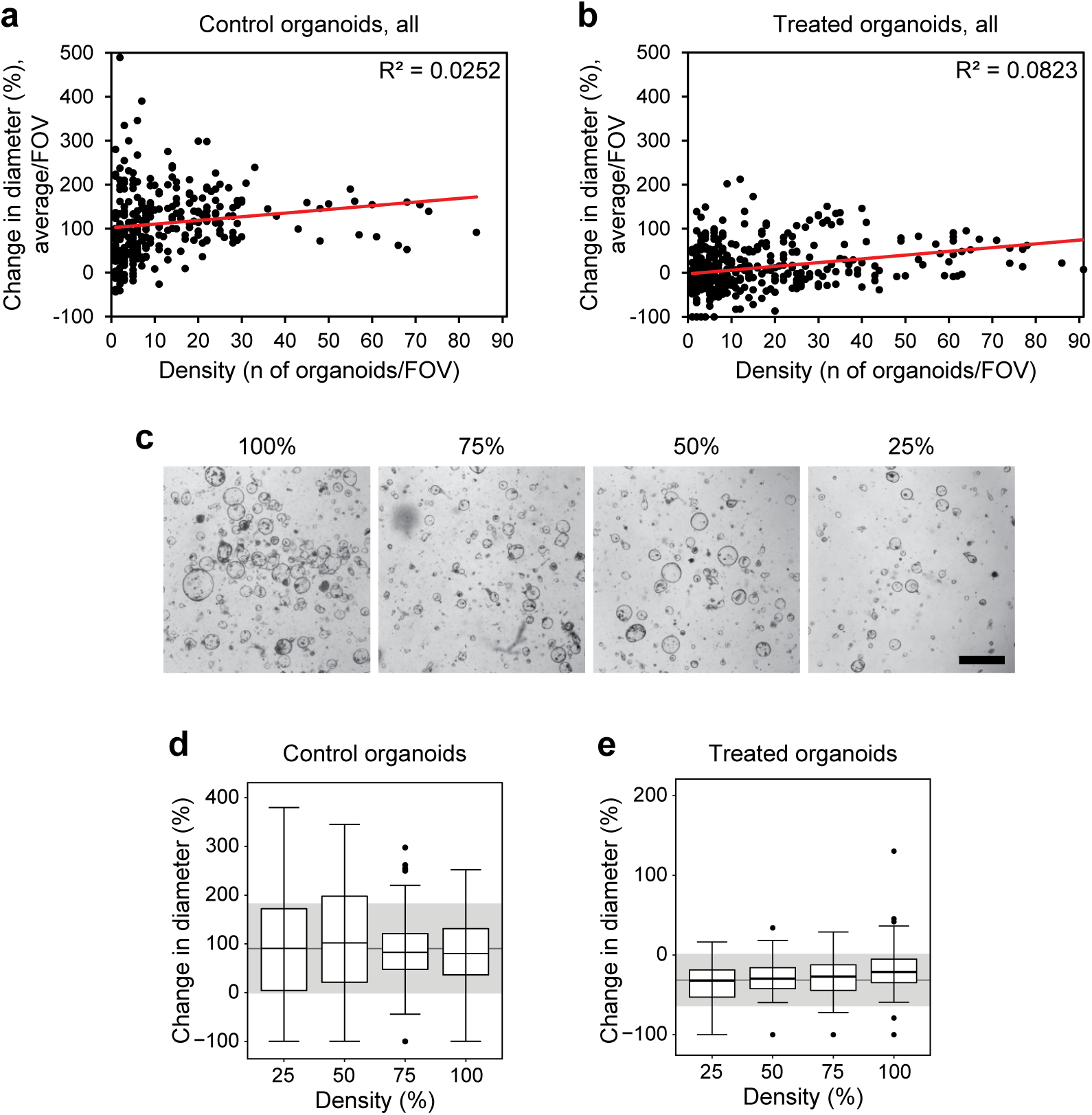
Density of the MDCOs in the Matrigel matrix does not affect growth or drug response. Scatter plots display if the density of MDCO cultures affect (a) growth or (b) drug response (n = 3580 and n = 6170, respectively). Density was determined by calculating the number of organoids per field of view (FOV) and was plotted against the average change in diameter per FOV. Linear trend lines are indicated in red. The R2 values displayed in the top right corner indicate the lack of significant correlation between change in diameter and the culture density. (c) Representative images illustrating starting density percentages used for the prospective analyses to validate observations seen in (a) and (b). Box and whisker plots displaying (d) control organoids and (e) treated organoids at different culture densities in the prospective study (n = 362 and n = 498). In all box and whisker plots the grey line indicates the population change in diameter mean while the grey shading indicates one standard deviation above and below the population mean. Representative images from (c) are shown at the same magnification. Size bar, 1 mm.

This observation was confirmed in a prospective study where APPK MDCOs were plated at densities ranging from 25%-100% (Fig 6c) and allowed to mature for 24 hours. Baseline 4x brightfield images were taken and the overlayed media was exchanged with new media containing copanlisib (200nmol/L) or control. After a 48 hour incubation, the same MDCOs from day 0 were imaged again. The changes in diameter of control or treated MDCOs were compared across the different densities. We observed no significant variation in the growth of the MDCOs due to the density as the majority of the MDCOs were within 1 SD of the PM (91 ± 92%). (Fig 6d) This was also observed in the copanlisib treated MDCOs (−31 ± 32.8%). (Fig 6e)

### Multivariate analysis confirms baseline culture conditions do not affect growth or therapeutic response

Finally, using a multivariate analysis, we confirmed that these baseline culture conditions do not cause unique clustering of organoids. A uniform manifold approximation and projection (UMAP) was used to examine four key variables for each individual organoid: baseline diameter, distance to the edge of the Matrigel droplet, day 2 diameter, and percent change in diameter over 48 hours. This dimension reduction technique is similar to *t-*distributed stochastic neighbor embedding (*t*-SNE) and principal component analysis (PCA) but is faster and able to preserve more of the global data structure than other dimensional reduction techniques [46]. UMAP representations showed that no clear clustering of organoids occurred due to MDCOs being derived from different mice of the same genotype (Fig 7a), location of the original tumor (Fig 7b), or passage number of MDCOs (Fig 7c). The only variable that showed clear clustering of MDCOs was whether the MDCO was treated with control media or a PI3K pathway inhibitor (Fig 7d). This analysis was also done with the CPA applied. Similar results were found demonstrating that different mice (Suppl fig 5a), location of the original tumor (Suppl Fig 5b), or passage number (Suppl Fig 5c) do not cause clear clustering of organoids. Only the treatment status of an organoid, i.e., if it was a control or treated organoid, showed clear clustering (Suppl Fig 5d).

**Figure 7.**
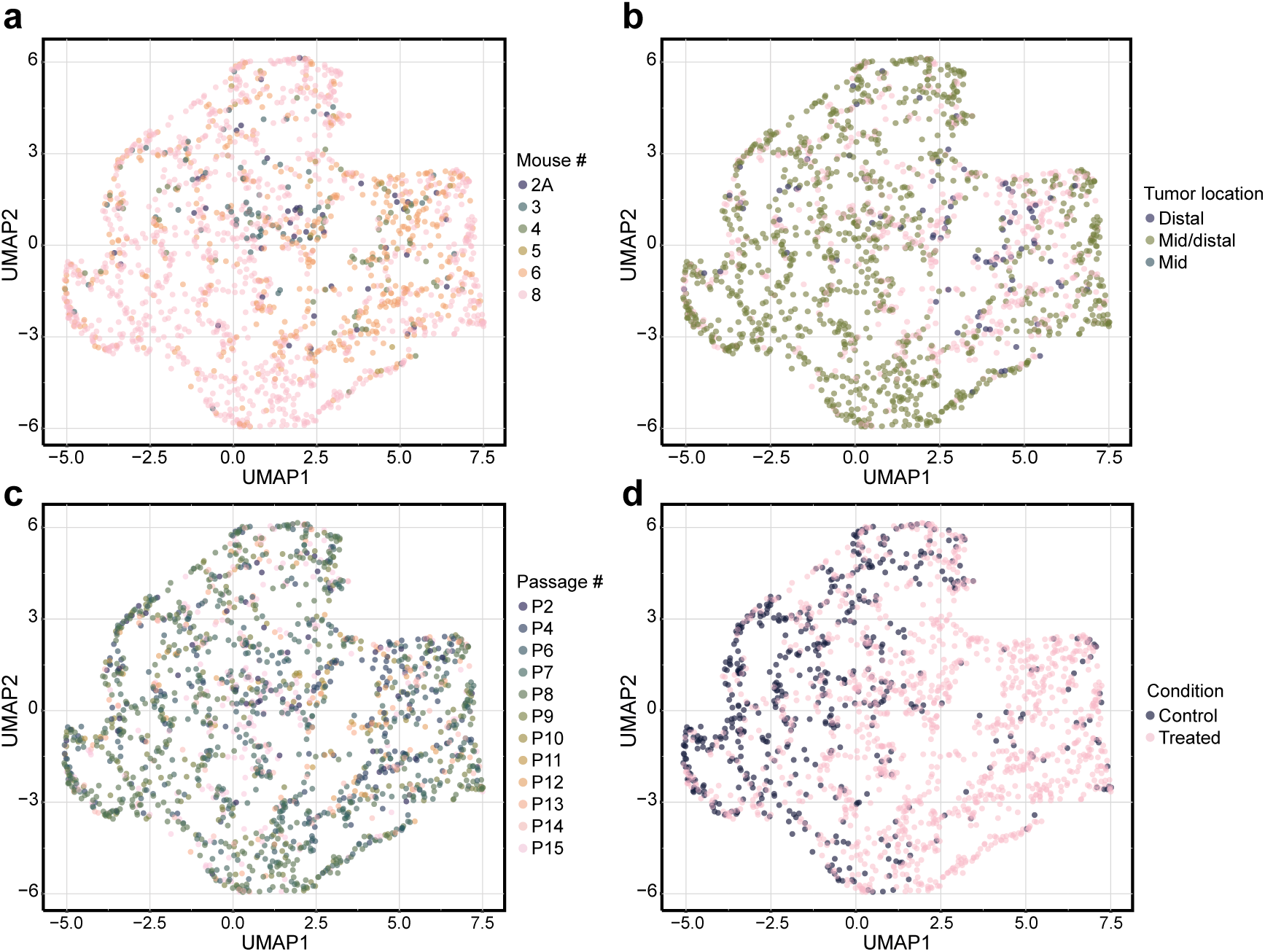
Multivariate analysis validates the finding that most baseline culture conditions do not affect growth or drug response. UMAP data-reduction examined four key variables (baseline diameter, distance to the edge of the Matrigel droplet, day 2 diameter, and percent change in diameter) to determine if baseline culture conditions caused clustering of individual MDCOs. UMAP visual representations showed that (a) MDCOs derived from different mice of the same genotype, (b) location of the original tumor within the colon, or (c) passage number did not cause clustering of MDCOs. Only (d) control and treated MDCOs showed any clear separation. (n = 1835 organoids)

## Discussion

Three-dimensional (3D) cancer organoids, derived from patient or mouse cancers, are becoming an increasingly popular model to identify and study novel combination therapies across many cancer types. Our group and others have shown that these *in vitro* models better recapitulate the genetic, morphological, and phenotypic characteristics of the cancers from which they were derived compared to traditional 2D immortalized cell lines [8-21].

In many cancer types, including CRC, there is a growing appreciation for how the molecular profile of a tumor will affect therapeutic response [3]. Therefore, patient-derived cancer organoids provide an attractive model to predict patient response and guide clinical treatment decisions [10, 19, 21, 29, 30, 33]. Alternatively, MDCOs are readily isolated from various transgenic mouse models across mutational profiles and diseases, providing a more controlled and high-throughput platform for screening potential therapies.

Given the recent development of cancer organoids, there have yet to be well-validated methods to determine response in these 3-dimentional cultures. Traditional 2D culture therapeutic response assessment techniques do not readily transfer to 3D organoid cultures as even basic culturing methods differ between 2D and 3D cultures. Many groups use methods that assess response on a well-by-well basis. Additionally, many assay readouts, such as CellTiterGlo, rely on the addition of reagents that do not allow for longitudinal assessment [9, 14, 16, 47-53]. These types of assays ignore many of the baseline characteristics of the organoids including heterogeneity within a culture. Some groups, including ours, evaluate the organoids on an individual basis using change in diameter or volume [10, 29-32]. Not only is this a less invasive method to determine response, but individual organoids can be tracked repeatedly throughout a longer study. This individual organoid level assessment also allows us to capture subpopulations and heterogeneity within populations (Fig 1).

If these organoid cultures are to be used to identify potential novel therapeutic strategies, and the methods of assessment standardized, it is important to understand how the culture conditions might affect the assays. Previously our group has shown that MTORC1/2 inhibition is necessary for a response in *Apc* and *Pik3ca* mutant CRC using the APPK MDCOs [40]. For this reason, we used this model in the context of PI3K pathway inhibition to determine if various culture conditions affect response. Notably, our MDCO lines and corresponding data were collected over the course over a 5-year period and from mice established across many generations. This allowed us to confirm that direct comparisons of data collected years apart from different cultures could be made.

We had previously observed and reported with a smaller cohort of MDCOs that those that were larger in size tended to grow at a slower rate [40]. With this larger cohort, which included those MDCOs originally assessed, we determined that APPK MDCOs ≥308 µm in diameter at baseline have a slower growth rate. We found that approximately 12% of the MDCOs analyzed here were ≥308 µm in diameter at the start of their study (Fig 2). These were excluded and all the mouse and culture conditions were reanalyzed. Excitingly, many of the results were better corroborated with the exclusion of the larger MDCOs as illustrated by more MDCOs growth and response rates within 1 standard deviation of the population mean and smaller R^2^ values (Suppl Figs 1-5).

We then broadly examined if any variation was seen between cultures derived from different mice of the same genotype or tumors isolated from different regions of the large intestine. While all mice were the same genotype, it is important to determine if any significant variation was seen among different mice across generations or even from different regions within the large intestine as gross histology varies slightly throughout the intestine. Significant variation between different mice or different regions of the colon could indicate some underlying differences in the biology of these mice or large intestine. For example, the gross histology of the proximal colon differs from the mid and distal colon in that it displays a herringbone pattern while the mid and distal colon is relatively smooth. Control and treated MDCOs isolated from 8 different mice and control MDCOs from different regions of the large intestine showed no significant difference in growth or response. However, some variation in response was seen in those treated MDCOs isolated from the proximal colon when compared to those from the other locations. The gross histological differences may account for some of the variation seen in these treated MDCOs compared to the mid, mid/distal, distal tumors (Fig 3). Altogether this data demonstrates that comparisons of growth and response can be made between MDCOs derived from different mice of the same genotype and from tumors from different regions of the large intestine which allows for direct comparison across generations of new litters.

Once we established that growth and response of MDCOs can be reliably examined across generations of mice, we assessed the influence of MDCO culture conditions. It is well established that immortalized 2-dimentional cultures can have significant changes as they increase in passage number due to genetic and phenotypic drift. We confirmed that no significant difference in growth or response develop as MDCOs are maintained in culture, up to passage 15 (Fig 4). Further studies are needed to assess whether culture beyond 15 passages promotes drift in MDCOs.

A unique aspect of MDCO cultures is their growth and maintenance in an extracellular matrix such as Matrigel. These unique culture conditions may provide a physical challenge for nutrient and drug delivery to MDCOs in the centermost region of the Matrigel droplet. We sought to address this concern by measuring MDCOs’ location within the Matrigel droplet to compare the growth and response between MDCOs at the center and edge of the droplet. We evaluated MDCOs as far as 4mm deep and found no correlation between MDCO location within the droplet and growth or response to PI3K pathway inhibitors. This demonstrates that the Matrigel droplet does not pose a significant barrier for nutrient and drug distribution (Fig 5).

Arguably the most important feature of organoid cultures is their ability to grow as 3D organized structures. When traditional immortalized 2D cultures are grown as 3D structures, they are usually pelleted cells and not organized in a coherent fashion. This is potentially due to the result of two key but opposing features that 2D cell lines can possess: contact inhibition and overgrowth. In both cases, proliferation rates are significantly altered with confluency, which can affect therapeutic response [4-6, 43-45]. We observed that neither growth nor response was altered due to the density of the culture (number of organoids/FOV). This was prospectively confirmed with an MDCO line cultured to its greatest density and diluted down to 25% of its highest density illustrating that MDCOs at different densities can be compared (Fig 6). Additionally, this demonstrates that MDCO cultures are more cancer-like, compared to many 2D cultures, in their ability to propagate without contact inhibition.

Finally using the dimension reduction technique UMAP, we demonstrated that MDCOs only separate based on whether they were treated with an inhibitor or control media. This multivariate analysis simultaneously evaluated many of these baseline culture conditions to corroborate the finding that MDCOs from different mice of the same genotype, or tumors from different locations, or passage number do not affect growth or response in these APPK MDCOs (Fig 7).

Further work should determine if these results hold true in other newly established organoid models of other diseases. Altogether we demonstrate that MDCO lines isolated from different mice of the same genotype and different tumors within the large intestine across several years of mouse colony propagation can be directly compared. Importantly, most of the culture conditions, including passage number, density, and location within the Matrigel droplet do not influence how an individual MDCO will grow or respond to a given therapy. We did confirm, however, that the starting size of an MDCO can influence its growth rate and subsequent response to therapy. Additionally, our work demonstrates the importance of examining response at an individual organoid level as opposed to at the population or well level. These studies bring to light the importance of understanding potentially confounding features to any assay, particularly drug assessment in 3D organoid models.

## Methods

### Cell isolation and organoid culture

All animal studies were performed adhering to approved protocols by the Institutional Animal Care and Use Committee at the University of Wisconsin (Madison, WI) following the guidelines of the American Association for the Assessment and Accreditation of Laboratory Animal Care International. *Apc*^*fl/fl*^ mice (B6.Cg-*Apc*^*tm2Rak*^; NCL Mouse Repository; Strain number 01xAA), *Pik3ca*^*H047R*^ mice (FVB.129S6 Gt(ROSA)26Sortm1(Pik3ca*H1047R)Egan/J; The Jackson Laboratory; Stock Number 016977) and *Fc*^*2*^ mice [FVB/N-Tg(Fapb1-Cre)1Jig; NCI Mouse Repository; Strain number 01XD8] were used to generate *Fc*^*1*^*Apc*^*fl/+*^*Pik3ca*^*H1047R*^ mice and these mice were genotyped as previously described [54, 55].

Colorectal cancer cells were isolated from *Fc*^*1*^ *Apc*^*fl/+*^ *Pik3ca*^*H1047R*^ (APPK) mice and cultured in Matrigel (Corning, cat #75796-276) as previously described [42].

### Pharmacologic agents

The PI3K pathway inhibitors AZD2014 (HY-15247, MedChem Express), everolimus (E-4040, LC Laboratories), NVP-BEZ235 (N-4288, LC Laboratories), TAK228 (I-3344, LC Laboratories), and BYL-719 (CT-BYL719, ChemiTek) were dissolved in DMSO to a 10mmol/L stock concentration. Copanlisib (HY-15346, MedChem Express) was dissolved in 10mmol/L TFA/DMSO to a 5mmol/L stock concentration. Described inhibitors were diluted in fresh media at concentrations ranging from 5nmol/L to 500nmol/L for diameter studies.

### Baseline culture conditions analysis

Organoids were plated in 24-well tissue culture plates and allowed to mature for 24-96 hours. Passages between 1 and 15 were used for the described studies. Prior to treatment, baseline 4x images were taken on a Nikon Ti-S inverted microscope. After imaging, overlayed feeding media was replaced with fresh feeding media containing drug. Following 48 hours of incubation at 37°C and 5% CO_2_, posttreatment images were taken. All images were analyzed using ImageJ, measuring the longest diameter of each organoid and the shortest distance from the organoid edge to the Matrigel edge. In each study, metrics of baseline organoid size, passage number (1-15), plating density, location within the Matrigel droplet, and the mouse identification number were collected for analysis.

For studies examining differences in dilutions, APPK MDCOs were cultured to their highest density and diluted to 75%, 50%, and 25% of the highest density. Cell suspensions were then plated in a 1:1 ratio with Matrigel as previously described. Response was assessed by exchanging feeding media with fresh media containing copanlisib (200nmol/L) or control and diameter studies continued as previously described [40].

### UMAP Analysis

Clustering of organoids across all mice and treatment conditions was represented using Uniform Manifold Approximation and Projection (UMAP) with the UMAP package in Python v3.7 [46]. UMAP dimensionality reduction was performed on four key variables (Day 0 Diameter, Distance to Edge, Day 2 Diameter, Percent Change in Diameter) for projection in 2D space. Organoids without one of these measurements were removed from the analysis, which left 1835 organoids included in the UMAP. The following parameters were used for UMAP visualizations: “n _neighbors”: 200; “min_dist”: 0.2, “metric”: cosine, “n_components”: 2. The generated UMAP data frame was then merged with the original input data frame for visualization in R (R-studio v.1.4).

### Statistical analysis

A change point analysis is a statistical analysis used to determine if a change in ordered data has occurred. A change point analysis was conducted to assess the impact of baseline size [56]. The percent change in organoid diameter over 48 hours was used to analyze growth and treatment response across baseline culture conditions. Kernel density plots using the *sm* package, Tukey’s boxplots, and scatter plots were generated using the *ggplot2* package and R software [57-59]. Glass’s delta was used to calculated treatment effect size [41].

## Author contributions

R.A. DeStefanis and A. Olson contributed equally.

R.A. DeStefanis, J.D. Kratz, and D.A. Deming conceived and designed the study. R.A. DeStefanis, A. Olson, J.D. Kratz, and D.A. Deming developed the methodology. R.A. DeStefanis, A. Olson, G. Sha, A DeZeeuw, A.A. Gillette, and K.A Johnson acquired the data and did the analysis and/or interpretation. All authors wrote, reviewed and/or revised the manuscript. L.Clipson, and C.A. Pasch provided administrative, technical and/or material support. M.C. Skala and D.A. Deming were the study supervisors.

## Additional information

Dr. Deming has received research support from Bayer and Takeda. He has also been compensated for serving on a scientific advisory board for Bayer. The other authors declare no potential conflict of interest.

## Notes

**Funding/Acknowledgements** This project was supported by NIH grants R37 CA226526 (DAD), T32 AG000213 (JDK), T32 CA009135 (KAJ), and P30 CA014520 (Core Grant, University of Wisconsin Carbone Cancer Center). The Skala laboratory is supported by NIH grants R01 CA185747 (MCS), R01 CA205101 (MCS), R01 CA211082 (MCS), U01 HL145792 (MCS). Additional support was provided from Funk Out Cancer (DAD), the Cathy Wingert Colorectal Cancer Research Fund (DAD), and the ACI/Schwenn Family Professorship (DAD).

## References

1. American Cancer Society. Key statistics for colorectal cancer. https://www.cancer.org/cancer/colon-rectal-cancer/about/key-statistics.html Accessed 8/19/2021.

2. Fight Colorectal Cancer. https://fightcolorectalcancer.org/about-colorectal-cancer/general-information/facts-stats/ Accessed 8/19/2021.

3. Benson, A. B. et al. Colon Cancer, Version 2.2021, NCCN Clinical Practice Guidelines in Oncology. J Natl Compr Canc Netw 19, 329–359 (2021).

4. Tveit, K. M., & Pihl, A. Do cell lines in vitro reflect the properties of the tumours of origin? A study of lines derived from human melanoma xenografts. Br J Cancer 44, 775–786 (1981).

5. Esquenet, M., Swinnen, J. V., Heyns, W., & Verhoeven, G. LNCaP prostatic adenocarcinoma cells derived from low and high passage numbers display divergent responses not only to androgens but also to retinoids. J Steroid Biochem Mol Biol 62, 391–399 (1997).

6. Daniel, V. C. et al. A primary xenograft model of small-cell lung cancer reveals irreversible changes in gene expression imposed by culture in vitro. Cancer Res 69, 3364–3373 (2009).

7. Xue, X., & Shah, Y. M. In vitro organoid culture of primary mouse colon tumors. J Vis Exp, e50210 (2013).

8. van de Wetering, M. et al. Prospective derivation of a living organoid biobank of colorectal cancer patients. Cell 161, 933–945 (2015).

9. Pauli, C. et al. Personalized In Vitro and In Vivo Cancer Models to Guide Precision Medicine. Cancer Discov 7, 462–477 (2017).

10. Pasch, C. A. et al. Patient-Derived Cancer Organoid Cultures to Predict Sensitivity to Chemotherapy and Radiation. Clin Cancer Res 25, 5376–5387 (2019).

11. Vlachogiannis, G. et al. Patient-derived organoids model treatment response of metastatic gastrointestinal cancers. Science 359, 920–926 (2018).

12. Ashley, N., Jones, M., Ouaret, D., Wilding, J., & Bodmer, W. F. Rapidly derived colorectal cancer cultures recapitulate parental cancer characteristics and enable personalized therapeutic assays. J Pathol 234, 34–45 (2014).

13. Dijkstra, K. K. et al. Generation of Tumor-Reactive T Cells by Co-culture of Peripheral Blood Lymphocytes and Tumor Organoids. Cell 174, 1586–1598 e1512 (2018).

14. Ganesh, K. et al. A rectal cancer organoid platform to study individual responses to chemoradiation. Nat Med 25, 1607–1614 (2019).

15. Kondo, J. et al. Retaining cell-cell contact enables preparation and culture of spheroids composed of pure primary cancer cells from colorectal cancer. Proc Natl Acad Sci U S A 108, 6235–6240 (2011).

16. Schutte, M. et al. Molecular dissection of colorectal cancer in pre-clinical models identifies biomarkers predicting sensitivity to EGFR inhibitors. Nat Commun 8, 14262 (2017).

17. Schumacher, D. et al. Heterogeneous pathway activation and drug response modelled in colorectal-tumor-derived 3D cultures. PLoS Genet 15, e1008076 (2019).

18. Johnson, K. A. et al. Human Colon Organoids and Other Laboratory Strategies to Enhance Patient Treatment Selection. Curr Treat Options Oncol 21, 35 (2020).

19. Sharick, J. T. et al. Cellular Metabolic Heterogeneity In Vivo Is Recapitulated in Tumor Organoids. Neoplasia 21, 615–626 (2019).

20. Weeber, F. et al. Preserved genetic diversity in organoids cultured from biopsies of human colorectal cancer metastases. Proc Natl Acad Sci U S A 112, 13308–13311 (2015).

21. Arnadottir, S. S. et al. Characterization of genetic intratumor heterogeneity in colorectal cancer and matching patient-derived spheroid cultures. Mol Oncol 12, 132–147 (2018).

22. Dangles-Marie, V. et al. Establishment of human colon cancer cell lines from fresh tumors versus xenografts: comparison of success rate and cell line features. Cancer Res 67, 398–407 (2007).

23. Mullins, C. S. et al. Integrated Biobanking and Tumor Model Establishment of Human Colorectal Carcinoma Provides Excellent Tools for Preclinical Research. Cancers (Basel) 11, (2019).

24. Boehnke, K. et al. Assay Establishment and Validation of a High-Throughput Screening Platform for Three-Dimensional Patient-Derived Colon Cancer Organoid Cultures. J Biomol Screen 21, 931–941 (2016).

25. Phan, N. et al. A simple high-throughput approach identifies actionable drug sensitivities in patient-derived tumor organoids. Commun Biol 2, 78 (2019).

26. Boretto, M. et al. Patient-derived organoids from endometrial disease capture clinical heterogeneity and are amenable to drug screening. Nat Cell Biol 21, 1041–1051 (2019).

27. Kopper, O. et al. An organoid platform for ovarian cancer captures intra-and interpatient heterogeneity. Nat Med 25, 838–849 (2019).

28. Ponz-Sarvise, M. et al. Identification of Resistance Pathways Specific to Malignancy Using Organoid Models of Pancreatic Cancer. Clin Cancer Res 25, 6742–6755 (2019).

29. Sunil, A. et al. Abstract 1494: Etiologies of patient-derived colorectal cancer organoid growth heterogeneity across multiple patient samples and culture conditions. Cancer Research 80, 1494–1494 (2020).

30. Gil, D. A., Deming, D., & Skala, M. C. Patient-derived cancer organoid tracking with wide-field one-photon redox imaging to assess treatment response. J Biomed Opt 26, (2021).

31. Zoetemelk, M., Rausch, M., Colin, D. J., Dormond, O., & Nowak-Sliwinska, P. Short-term 3D culture systems of various complexity for treatment optimization of colorectal carcinoma. Sci Rep 9, 7103 (2019).

32. Kaushik, G. et al. Selective inhibition of stemness through EGFR/FOXA2/SOX9 axis reduces pancreatic cancer metastasis. Oncogene 40, 848–862 (2021).

33. Sharick, J. T. et al. Metabolic Heterogeneity in Patient Tumor-Derived Organoids by Primary Site and Drug Treatment. Front Oncol 10, 553 (2020).

34. Xie, B. Y., & Wu, A. W. Organoid Culture of Isolated Cells from Patient-derived Tissues with Colorectal Cancer. Chin Med J (Engl) 129, 2469–2475 (2016).

35. Boj, S. F. et al. Organoid models of human and mouse ductal pancreatic cancer. Cell 160, 324–338 (2015).

36. Yin, X. et al. Niche-independent high-purity cultures of Lgr5+ intestinal stem cells and their progeny. Nat Methods 11, 106–112 (2014).

37. Sato, T. et al. Long-term expansion of epithelial organoids from human colon, adenoma, adenocarcinoma, and Barrett’s epithelium. Gastroenterology 141, 1762–1772 (2011).

38. Tao, Y. et al. Aging-like Spontaneous Epigenetic Silencing Facilitates Wnt Activation, Stemness, and Braf(V600E)-Induced Tumorigenesis. Cancer Cell 35, 315–328 e316 (2019).

39. Nam, M. O. et al. Effects of a small molecule R-spondin-1 substitute RS-246204 on a mouse intestinal organoid culture. Oncotarget 9, 6356–6368 (2018).

40. Fricke, S. L. et al. MTORC1/2 Inhibition as a Therapeutic Strategy for PIK3CA Mutant Cancers. Mol Cancer Ther 18, 346–355 (2019).

41. Glass, G. V., McGraw, B., & Smith, M. L. Meta-analysis in social research. London: Sage. (1981).

42. Foley, T. M. et al. Dual PI3K/mTOR Inhibition in Colorectal Cancers with APC and PIK3CA Mutations. Mol Cancer Res 15, 317–327 (2017).

43. Sambuy, Y. et al. The Caco-2 cell line as a model of the intestinal barrier: influence of cell and culture-related factors on Caco-2 cell functional characteristics. Cell Biol Toxicol 21, 1–26 (2005).

44. Chang-Liu, C. M., & Woloschak, G. E. Effect of passage number on cellular response to DNA-damaging agents: cell survival and gene expression. Cancer Lett 113, 77–86 (1997).

45. Wilding, J. L., & Bodmer, W. F. Cancer cell lines for drug discovery and development. Cancer Res 74, 2377–2384 (2014).

46. McInnes, L., Healy, J., & Melville, J. UMAP: Uniform Manifold Approximation and Projection for Dimension Reduction. 1802.03426v3. https://arxiv.org/abs/1802.03426. (2018).

47. Driehuis, E., Kretzschmar, K., & Clevers, H. Establishment of patient-derived cancer organoids for drug-screening applications. Nat Protoc 15, 3380–3409 (2020).

48. Broutier, L. et al. Human primary liver cancer-derived organoid cultures for disease modeling and drug screening. Nat Med 23, 1424–1435 (2017).

49. Ooft, S. N. et al. Patient-derived organoids can predict response to chemotherapy in metastatic colorectal cancer patients. Sci Transl Med 11, (2019).

50. Mullenders, J. et al. Mouse and human urothelial cancer organoids: A tool for bladder cancer research. Proc Natl Acad Sci U S A 116, 4567–4574 (2019).

51. Duarte, A. A. et al. BRCA-deficient mouse mammary tumor organoids to study cancer-drug resistance. Nat Methods 15, 134–140 (2018).

52. Lohmussaar, K. et al. Assessing the origin of high-grade serous ovarian cancer using CRISPR-modification of mouse organoids. Nat Commun 11, 2660 (2020).

53. Hai, J. et al. Generation of Genetically Engineered Mouse Lung Organoid Models for Squamous Cell Lung Cancers Allows for the Study of Combinatorial Immunotherapy. Clin Cancer Res 26, 3431–3442 (2020).

54. Hung, K. E. et al. Development of a mouse model for sporadic and metastatic colon tumors and its use in assessing drug treatment. Proc Natl Acad Sci U S A 107, 1565–1570 (2010).

55. Saam, J. R., & Gordon, J. I. Inducible gene knockouts in the small intestinal and colonic epithelium. J Biol Chem 274, 38071–38082 (1999).

56. Killick, R., & Eckley, I. A. changepoint: An R Package for Changepoint Analysis. https://www.jstatsoft.org/article/view/v058i03. Journal of Statistical Software 58, (2014).

57. Bowman, A. W., & Assalini, A. R package ‘sm’: nonparametric smoothing methods (version 2.2-5.4). http://www.stats.gla.ac.uk/~adrian/sm, http://azzalini.stat.unipd.it/Book_sm. (2014).

58. R Core Team. R: A language and environment for statistical computing. R Foundation for Statistical Computing, Vienna, Austria. https://www.R-project.org/. (2021).

59. Wickham, H. ggplot2: Elegant Graphics for Data Analysis. Springer-Verlag, New York. (2016).

