## Supplemental figures for "Impact of baseline culture conditions of mouse-derived cancer organoids when determining therapeutic response and tumor heterogeneity"

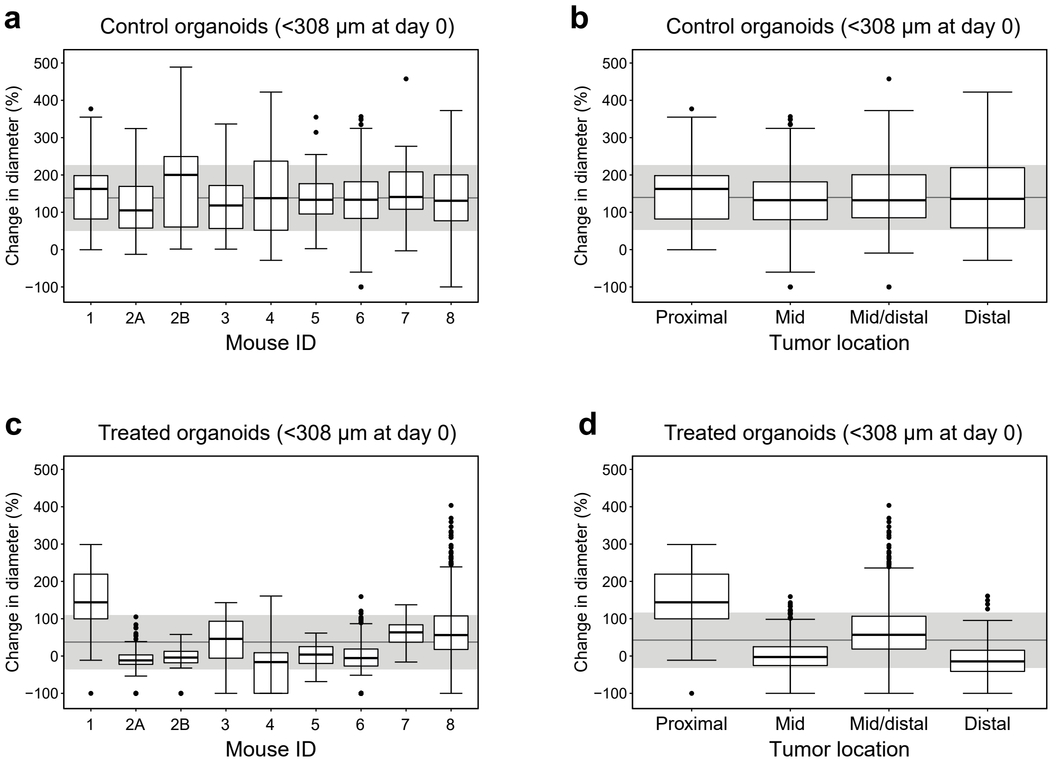


**Supplementary Figure S1. Effect of derivation of MDCOs from different mice of the same genotype or from different regions of the colon with change point analysis applied.** MDCOs with a starting diameter of ≥308 μm were removed from these analyses. Box and whisker plots displaying (a) control organoids and (c) treated organoids derived from different mice of the same genotype (n = 834 and n = 2222, respectively). Box and whisker plots displaying (b) control organoids and (d) treated organoids derived from tumors isolated from different regions of the large intestine (n = 734 and n = 2002, respectively). Note that the gross histology of the proximal colon is different from the mid and distal colon. For all box and whisker plots the grey line is the population mean with the grey shading indicating ±1 standard deviation from the mean.

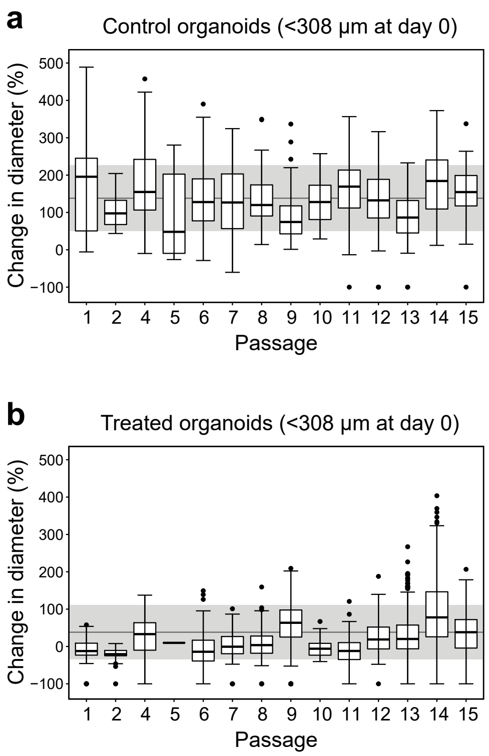


**Supplementary Figure S2. The passage number of MDCOs does not affect growth or drug response when change point is applied.** Box and whisker plots demonstrating (a) control organoids and (b) treated organoids at different passages (n = 817 and n = 2136 respectively). Passages range from 1-15 from isolation. The change point was applied and all MDCOs with a starting diameter of ≥308 µm were removed from these analyses. Note that passage 5 in the treated MDCOs only had n = 1. For all plots, the grey line is the population mean while the grey shading indicates ± 1 standard deviation from the mean.

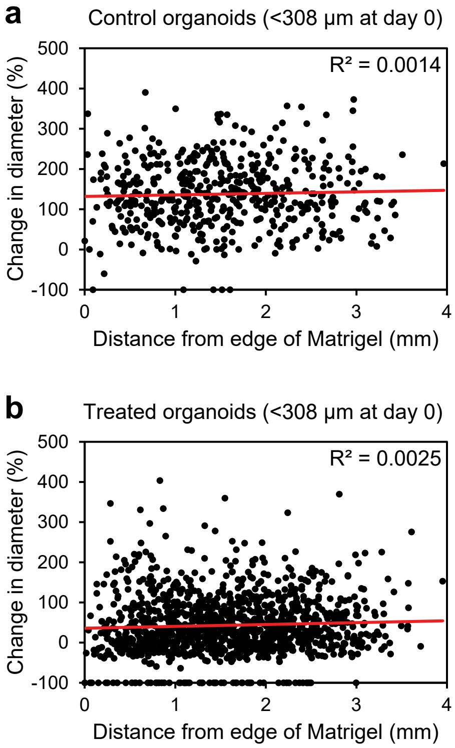


**Supplementary Figure S3. Location of the MDCOs within the Matrigel matrix does not affect growth or drug response with the change point applied.** Scatter plots display individual (a) control or (b) treated MDCOs distance from the edge of the Matrigel matrix (mm) plotted against its change in diameter (n = 493 and n = 1186, respectively). Change point analysis was applied and only MDCOs with a starting diameter size of <308 µm were included for analyses. The linear trend lines are indicated in red. Correlations were determined using R2 values displayed in the top right corner.

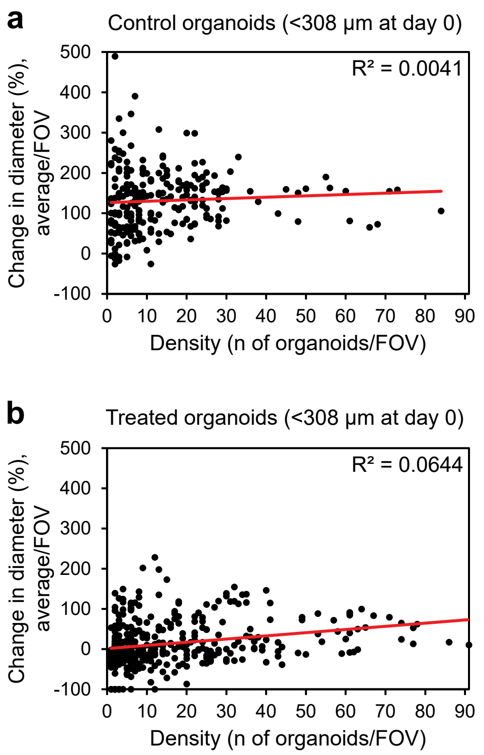


**Supplementary Figure S4. Density of the MDCOs does not affect growth or drug response with the change point applied.** Scatter plots demonstrate if the density of MDCO cultures affect (a) growth or (b) drug response (n = 3277 and n = 6021, respectively). The change point was applied and only the MDCOs with a starting diameter of <308μm were included for these analyses. For each plot, the average change in diameter per field of view (FOV) was plotted against the number of MDCOs per FOV. Linear trends for each plot are indicated in red. R2 values used to determine correlation are displayed in the top right corner of the plots.

**Suppl Figure S5a-b**

a

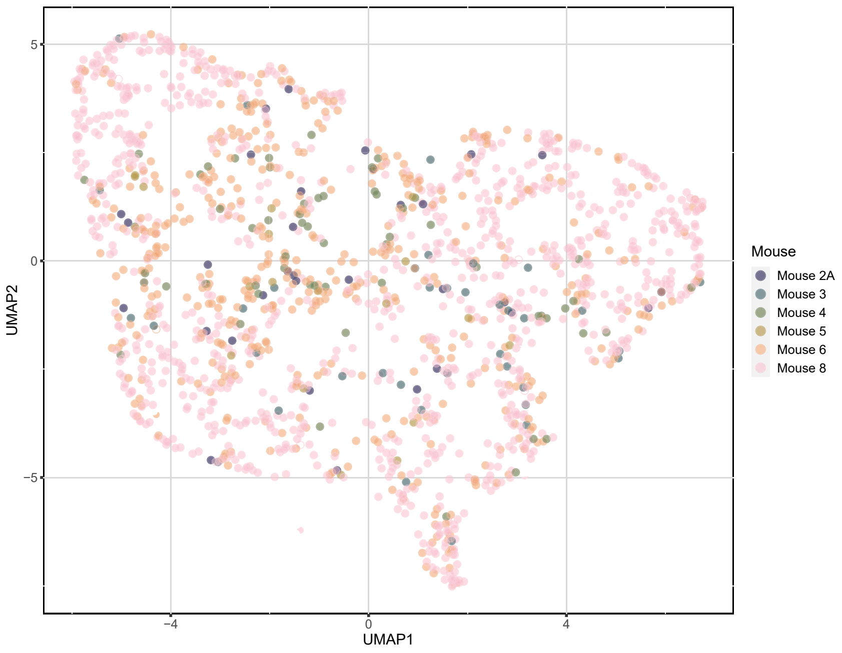


b

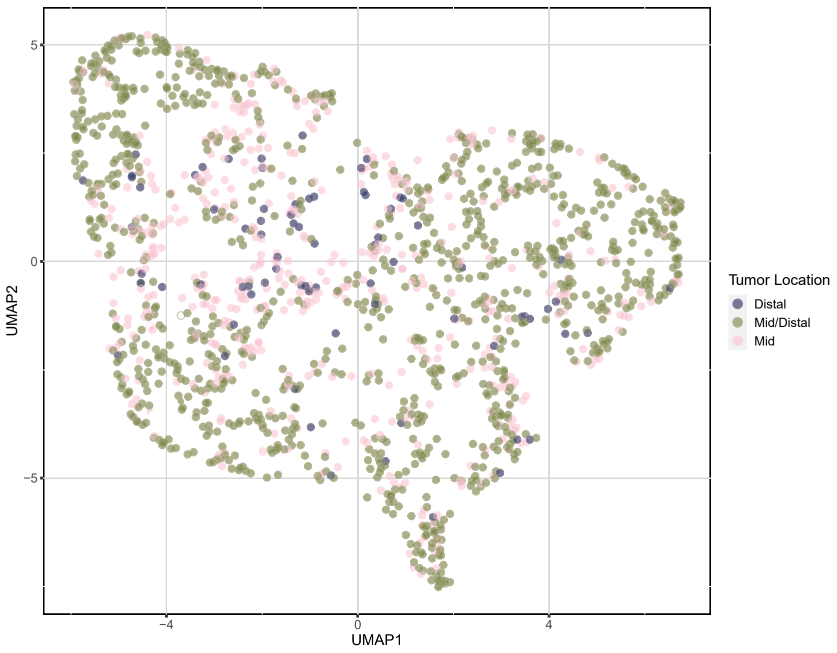

### Suppl Figure S5c-d

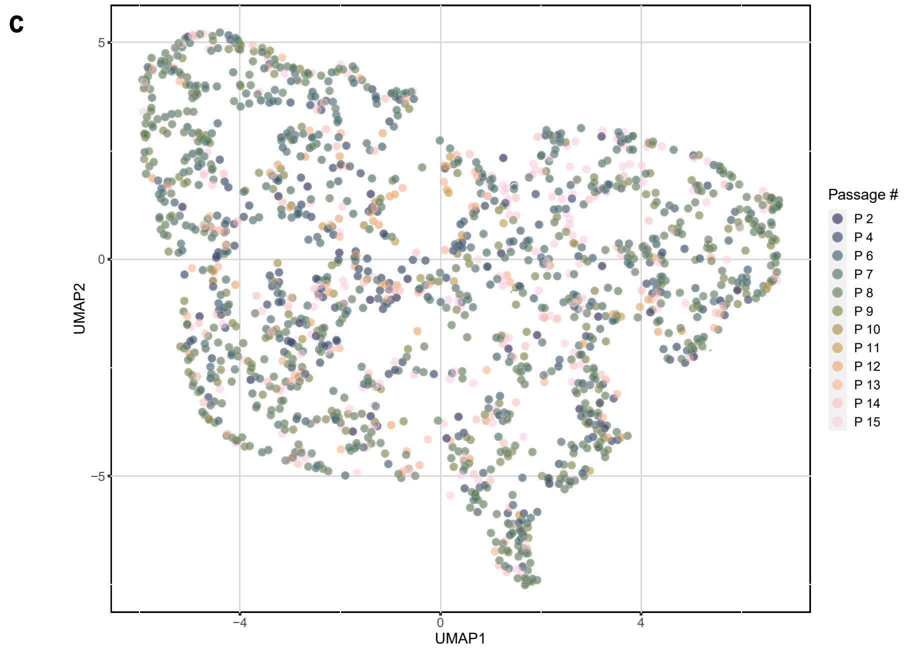


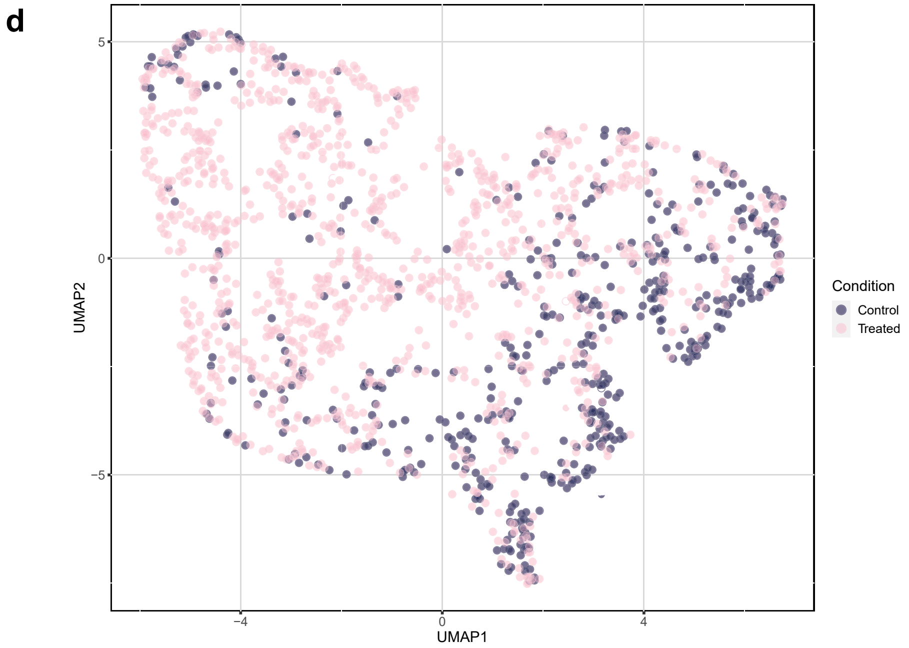


**Supplementary Figure S5. UMAP multivariate analyses with change point applied validates that most baseline culture conditions do not affect growth or drug response.** (a) Different mice of the same genotype, (b) location of original tumor within the colon, and (c) passage number did not cause any clustering of MDCOs. Only (d) treatment status of an MDCO showed any clustering. (n = 1614 organoids)
